# The transcriptome of acute dehydration in Myeloid Leukemia cells

**DOI:** 10.1101/2022.09.23.509183

**Authors:** David B. Mark Welch, Travis J. Gould, Ada L. Olins, Donald E. Olins

**Author notes:** Corresponding Author (DEO).

## Abstract

Live human myeloid leukemia (HL-60/S4) cells exposed to acute hyperosmotic stress with sucrose undergo dehydration and cell shrinkage. Interphase chromatin and mitotic chromosomes congeal, and exhibit altered phase separation (demixing) of chromatin-associated proteins. To investigate concurrent changes in the transcriptome, we exposed exponentially growing HL-60/S4 cells to acute hyperosmotic stress (∼600 milliOsmolar) for 30 and 60 minutes by addition of sucrose to the culture medium. We employed RNA-Seq of polyA mRNA to identify genes with significantly increased or decreased transcript levels relative to untreated control cells (i.e., differential gene expression). These identified genes were examined for over-representation of Gene Ontology (GO) terms. In hyperosmotically-stressed cells, multiple GO terms associated with transcription, translation, mitochondrial function and proteosome activity, as well as the gene set “replication-dependent histones”, were over-represented among genes with increased transcript levels; whereas, genes with decreased transcript levels were over-represented in various GO terms for transcription repressors. The overall transcriptome profiles of these stressed cells suggest a rapid acquisition of cellular rebuilding, a futile homeostatic response, as these cells are ultimately doomed to a dehydrated death.

“Do not go gentle into that good night

Rage, rage against the dying of the light”

Dylan Thomas (1914-1953)

## Introduction

Normal cells in our body are frequently exposed to hyperosmotic stress conditions [1]. These are generally brief challenges of resilient cell systems. More serious consequences can occur during tissue inflammation and various diseases [1]. While there has been considerable exploration of the adaptation of renal (kidney) cells to hyperosmotic stress, evidence suggests that other tissues respond differently [2] and there is a need to develop convenient reproducible models using other cell types. The influence of growth medium osmolarity upon in vivo mammalian tissue culture cell nuclear structure has been explored for several decades. Despite the variation of cell types studied, there is a consensus that acute hyperosmotic stress (e.g.,>300 milliOsmolar [mOsM], total) produces rapid cell volume shrinkage and heterogeneity of nuclear chromatin condensation [3-7]. In our previous study [7], we exposed live human myeloid leukemic HL-60/S4 cells to 300 mM sucrose in growth medium (i.e., ∼600 mOsM, total) producing acute dehydration and cell shrinkage. Employing microscopy, we observed interphase and mitotic chromatin condensation (denoted by us: “congelation”). The chromatin description “congelation” was first employed in [7]. This description is intended to distinguish hyperosmotic stress chromatin condensation from normal heterochromatin or mitotic chromosome condensation.

The effect on interphase nuclear chromatin can be clearly seen employing DAPI staining of DNA combined with STED imaging (Fig 1): the dispersed “fine” interphase nuclear chromatin fibers (in the absence of sucrose) “congeal” into thicker strands during hyperosmotic stress. Combined with immunofluorescent staining microscopy, we documented apparent phase separation with loss of colocalization of various chromatin-associated proteins (possible “demixing”) resulting from the acute hyperosmotic stress [7]. In this study, we did not focus on long-term incubation (i.e., > 1 hour) of HL-60/S4 cells in 300 mM sucrose, since the cells appear to deteriorate and die after several hours in these dehydrating conditions. In this regard, HL-60/S4 cells appear to differ from some other cell types that can adapt to ongoing hyperosmotic stress [8, 9].

**Fig 1.**
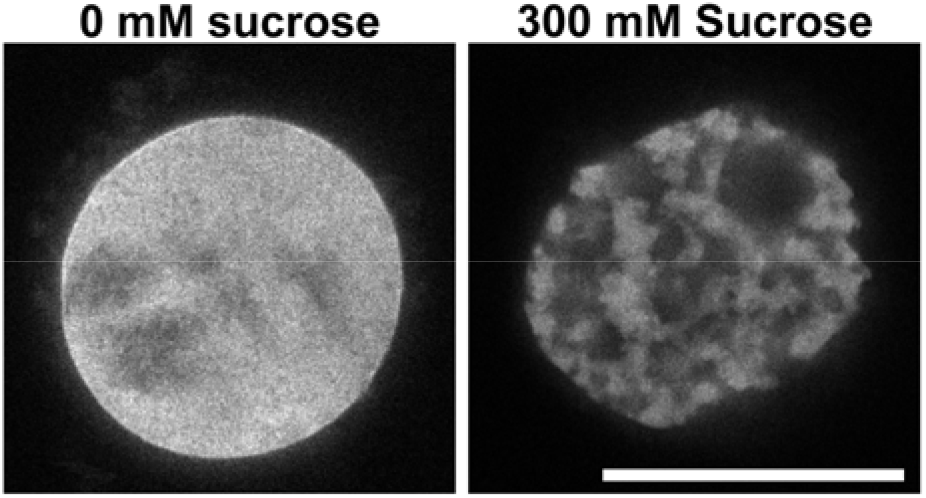
Effect of hyperosmotic stress upon interphase nuclear DNA distribution. STED images of undifferentiated HL-60/S4 cells after fixation, permeabilization and staining for DNA with DAPI. (Left Panel) Control cell nucleus, untreated in tissue culture medium. (Right Panel) Cell nucleus in tissue culture medium plus 300 mM sucrose for 30 minutes (∼600 mOsM). Magnification bar: 10 µm. Similar images can be found in Fig 4 of reference [7].

Two central questions of the present study are: 1) How is the polyA mRNA transcriptome (i.e., transcript levels) of cells exposed to acute hyperosmotic stress altered, in comparison to unstressed cells? 2) How can these altered transcript levels be interpreted to understand changes in the stressed cell physiological state? Two prior studies have been published toward this end: both employed microarray analyses of several hundred genes [10, 11]. Because these studies employed a different analysis method and examined different cell types, compared to the present study, a direct comparison is difficult. In the present study, we compared the transcriptomes of acute (i.e., 30 and 60 min.) hyperosmotically stressed undifferentiated HL-60/S4 cells to that of unstressed (control) cells. HL-60/S4 cells are derived from a human myeloid leukemia that preserves the ability to differentiate *in vitro* into stable cell states [12, 13]. We employed a standard RNA-Seq approach using polyA mRNA isolated from biological replicates of stressed and unstressed cells, and mapped reads back to the human genome as a proxy for the equilibrium mRNA transcript level of each gene at the time of sampling. Differential gene expression (DGE) analysis of ∼16,000 genes revealed statistically significant increases or decreases in transcript levels compared to control (unstressed) transcript levels. We then employed over-representation-analysis (ORA) to identify gene ontology (GO) terms enriched in genes with increased or decreased transcript levels. The resulting profiles of GO terms permit us to speculate on the probable physiological functions that are affected in the stressed cells (i.e., the altered cell physiological state).

## Materials and methods

### Cell culture

HL-60/S4 cells (ATCC CRL-3306) were cultivated in RPMI-1640 medium, plus 10% FCS and 1% Pen/Strep. Rapidly growing cells in medium were added to dry sucrose in T-25 flasks yielding 300 mM sucrose (∼600 mOsM, total) for 30 and 60 minutes, as described earlier [7]. Cell differentiation of HL-60/S4 into granulocytes and macrophage has also been described earlier [12].

### RNA purification

Quadruplicate samples of undifferentiated HL-60/S4 cells (5×10^6^ cells/sample) exposed to 0, 30, or 60 minutes of sucrose (total 12 samples) were centifuged, rapidly frozen and stored in liquid nitrogen (LN_2_). Samples were thawed by the addition of the RLT lysis buffer from the Qiagen RNeasy Mini Kit and RNA purified according to the manufacturer’s protocol. Care was taken to maintain RNAase-free conditions with RNAase Zap. RNA was eluted with molecular biology grade water, frozen in LN_2_ and shipped on dry ice to the Marshall University Genomics Core Facility. QC determinations (<u>all 12 samples</u> had a RIN score of 10), preparation of the libraries (Illumina mRNA library preparation Kit) and sequencing (Illumina HiSeq1500) was carried out at the Core Facility. The RNA-seq data created in this study is openly available at https://www.ncbi.nlm.nih.gov/bioproject/?term=PRJNA686972 in the NCBI Sequence Read Archive as BioProject accession PRJNA686972. Data and analyses of undifferentiated, retinoic acid (RA)-treated and phorbol ester (TPA)-treated HL-60/S4 cells can be found in a previous publication [12].

### Data analyses and presentation

We identified genes with significantly differential transcript levels following the RSEM-EBSeq workflow outlined at http://deweylab.github.io/RSEM and used the sequences and annotation of UCSC human genome v19 (hg19) from Illumina igenome: (https://support.illumina.com/sequencing/sequencing_software/igenome.html). Hg19 was chosen, instead of hg38, so that the hyperosmotic transcriptomes would be more comparable to the previous transcriptomes of undifferentiated and differentiated HL-60/S4 cells [12]. Furthermore, hg38 is a refinement of the assembly, including primarily alternatively spliced transcripts and a greater annotation of nonprotein-coding genes. Hg38 and the new hgnc names will recover more genes, but most of them are newly discovered and poorly annotated, so unlikely to have much impact on ORA and GO analyses. A bash script of the workflow is available at (Supporting Information) S8 Text. Briefly, we used bowtie2 v2.3.2 to map paired-end reads to transcripts extracted from the reference genome, and calculated transcript level values using RSEM v1.3.0. RSEM uses a maximum likelihood expectation-maximization algorithm to estimate the transcript levels of isoforms from RNA-Seq reads [14]. We then calculated the significance of relative expression differences using EBSeq v1.2.0 with the ng-vector option for isoform-level analysis [15]. EBSeq returns the normalized mean count of reads mapped using the median of ratios approach of DESeq [16] and the posterior probability of differential expression (PPDE) between control and treatment conditions, which is naturally corrected for multiple tests (i.e., the PPDE is equivalent to one minus the false discovery rate, FDR). We tested control versus 30 minutes and control versus 60 minutes of sucrose exposure, and accepted genes with PPDE>0.95 as having significantly different transcript levels. See S1 Table for a complete table of EBseq normalized mean counts and PPDE values for each hg19 gene. Output from RSEM and EBSeq were loaded into a MySQL database with hg19 annotation for analysis. We uploaded lists of differentially expressed genes (i.e., HGNC gene symbols) to WebGestalt (http://bioinfo.vanderbilt.edu/webgestalt) for over-representation analysis of GO non-redundant terms [17].

We considered a GO term over-represented if the hypergeometric test returned an adjusted p value (FDR) less than 0.05. See S2 Table for detailed output for all GO terms significantly over-represented after 30 minutes or 60 minutes of exposure to sucrose.

### Microscopy

STED imaging of DAPI stained HL-60/S4 cells +/-300 mM sucrose and Leica SP8 confocal imaging of HL-60/S4 cells +/-300 mM sucrose immunostained with rabbit anti-pPol II, rabbit anti-NPAT, mouse anti-epichromatin (PL2-6) and DAPI were performed by the same methods as described earlier [18]. Antibodies employed were HUABIO anti-Phospho RNA polymerase II POLR2A (S5) and Invitrogen (Thermo Fisher Scientific) anti-NPAT (PA5-668.39).

## Results

### Overall view of acute hyperosmotic stress effects on the mRNA transcriptome

Despite the considerable extent of interphase chromatin congelation at 30 and 60 minutes of hyperosmotic sucrose treatment, the relative transcript levels are essentially unchanged for most (∼60%) genes (Table 1 “Sucrose”). After 30 minutes of exposure, transcript levels are significantly increased for 3128 genes and significantly decreased for 2746. This change in transcript levels continues with ongoing exposure to sucrose: between 30 and 60 minutes the number of genes with significantly increased or decreased transcript levels changed by 103 and 215 genes, respectively. (See S1 Table for EBseq normalized mean counts and PPDE [statistical significance] values for each gene at a pairwise comparison of conditions.) The implication of these results is that most of the transcriptome changes occurred within the initial 30 minutes. It is useful to compare these hyperosmotic stress transcriptome changes to the transcriptome changes of chemically induced differentiation in HL-60/S4 cells for 4 days with retinoic acid (RA) into granulocytes or phorbol ester (TPA) into macrophage [12]. The differentiated cell states only exhibited a slightly greater change in relative transcript levels (Table 1 “Differentiation”). Observing cell differentiation in the microscope, (see Fig 1 in [13] and Fig 5 in [12]), indicates that during the 4 day induction period visible cellular phenotypes were exhibiting considerable changes. It is important to point out that the cells tested by hyperosmotic stress (this study) were <u>undifferentiated</u>. A lower % of genes remained unchanged after 4 days of differentiation, compared to the % of unchanged genes after 30 and 60 minutes of hyperosmotic sucrose stress.

**Table 1.** Significant Changes in Transcript Levels. Number of genes with significant changes in relative transcript levels for HL-60/S4 differentiation states [12] and for 30-and 60-minute exposure times to 300 mM sucrose (present study). Column titles: “DGE”, Differential Gene Expression; “RA”, 4 days of retinoic acid differentiation to granulocyte form; “TPA”, 4 days of phorbol ester differentiation to macrophage form; “30 and 60 min”, exposure time in medium+300 mM sucrose. “Control” is the total number of genes mapped under control conditions. “%” is the percentage of the appropriate “Total” for the specific cell condition. For a complete list of all analyzed 25,346 genes, see S1 Table.

| DGE | Differentiation |  |  |  | Sucrose |  |  |  |
| --- | --- | --- | --- | --- | --- | --- | --- | --- |
|  | RA | % | TPA | % | 30 min | % | 60 min | % |
| Increased | 4,249 | 26.2 | 5,156 | 31.2 | 3,128 | 18.4 | 3,231 | 20.3 |
| Decreased | 3,900 | 24.1 | 4,528 | 27.4 | 2,746 | 20.9 | 2,962 | 22.2 |
| Unchanged | 8,040 | 49.7 | 6,842 | 41.4 | 9,073 | 60.6 | 8,371 | 57.5 |
| Total | 16,189 |  | 16,526 |  | 14,947 |  | 14,564 |  |
| Control | 15,998 |  |  |  | 15,615 |  |  |  |

We performed over-representation-analysis of genes with increased and decreased transcript levels using each GO non-redundant annotation (i.e., Biological Processes, Cellular Component and Molecular Function). Analysis using genes with <u>increased</u> transcript levels revealed GO terms associated with transcription and translation, mitochondrial structure and function, and protein stress (Table 2; also, see S2 Table Biol., S3 Table Cell. and S4 Table Mol.) for complete ORA results with GO term identifiers, enrichment, and significance). The most enriched GO terms were associated with mitochondrial structure and function, which accounted for nearly 13% of all genes with increased transcript levels. In contrast, analysis using genes with <u>decreased</u> transcript levels yielded more diverse results, with GO terms reflecting a decrease in pathways involved in transcription repression and heterochromatin formation. Overall, these results suggest that the dehydrated physiological state of the stressed HL-60/S4 cells involves attempts to accomplish transcription and translation, and to increase mitochondrial oxidative phosphorylation, possibly to contend with protein misfolding and proteome degradation.

**Table 2.**
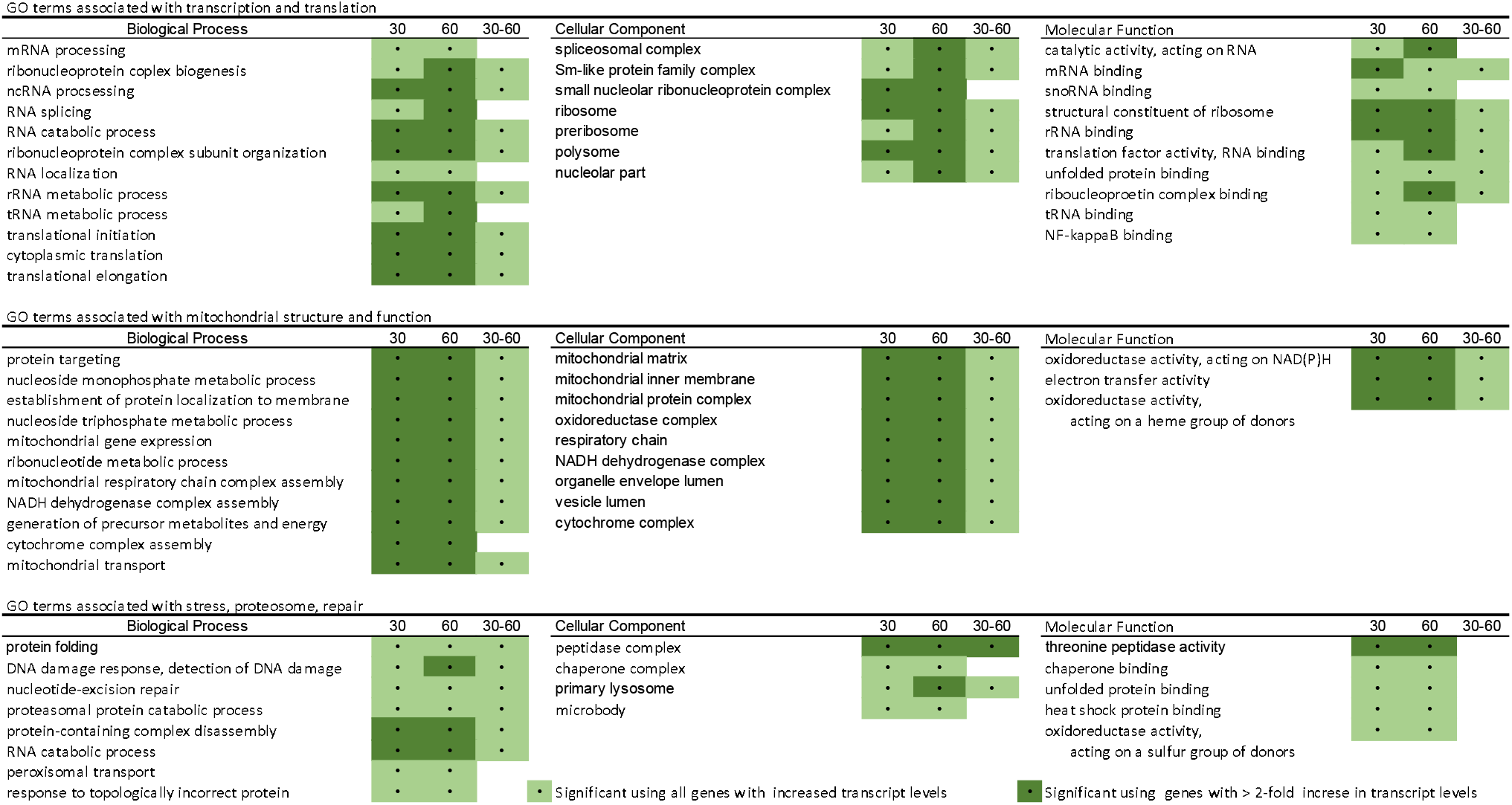
Summary of GO terms. Summary of GO terms from each functional domain which is enriched in genes with <u>increased relative transcript levels</u> in response to acute hyperosmotic stress, exposed for 0-30 min, 0-60 min and 30-60 min intervals. Color coding: light green, enriched among genes with where log2FC>0; dark green, enriched among genes where log2FC>1 (i.e., at least a doubling compared to control levels). See (Supporting Information) S2 Table Biol., S3 Table Cell. and S4 Table Mol for complete GO data in the three GO annotations (**a** Biological Processes; **b** Cellular Component; **c** Molecular Function). For definitions of GO terms and lists of included genes, see http://geneontology.org

### Nuclear distribution of “activated” RNA Polymerase II

Observing the structural transition of dehydrated interphase nuclear chromatin (Fig 1), we wondered how the spatial distribution of “activated” RNA Polymerase II (phosphorylated RNA Pol II) would be affected by hyperosmotic stress, relative to its distribution in untreated cells. Fig 2 displays confocal immunostaining images of anti-phosphorylated RNA Pol II, in relation to DAPI stained interphase nuclear chromatin (+/-300 mM sucrose for 30 minutes). These images clearly indicate that “activated” RNA Pol II persists within the interphase nuclei of hyperosmotically stressed cells. However, there is an apparent redistribution of “activated” Pol II staining to locations near the surface of congealed chromatin and within spaces between congealed chromatin.

**Fig 2.**
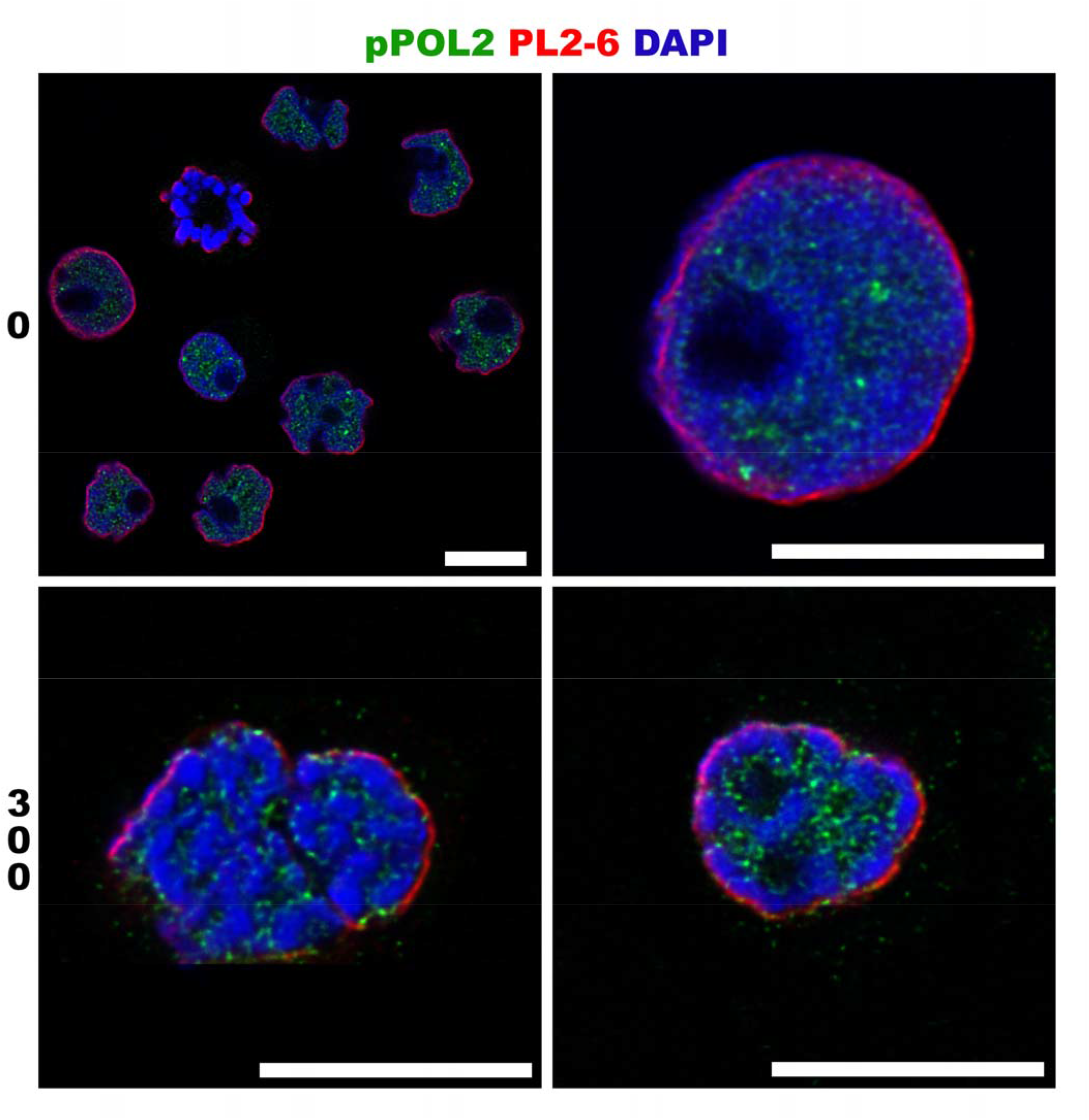
Effect of hyperosmotic stress upon interphase distribution of “activated” phosphorylated RNA Polymerase II. Confocal images of undifferentiated HL-60/S4 cells after formaldehyde fixation (+/-300 mM sucrose), permeabilization and immunostaining. (Top Row “0”) Control cells, untreated with sucrose in tissue culture medium. Note the absence of “activated” RNA Polymerase within mitotic chromosomes (yellow arrowhead), but presence within interphase nuclei. (Bottom Row “300” for a) Interphase cell nuclei in tissue culture medium containing 300 mM sucrose (∼600 mOsM) for 30 minutes prior to fixation. Staining: pPol II, phosphorylated (activated) RNA Polymerase II (green); PL2-6, anti-epichromatin, i.e., exposed nucleosome acidic patches at the surface of interphase chromatin (red), see references [7, 18, 19]; DAPI, DNA (blue). Magnification bar for all images: 10 µm.

### Selected examples of the transcriptome data

Given the extent and depth of the mRNA transcriptome data, we have chosen to discuss only a few prominent stress-upregulated cellular functions. The data are presented in two types of graphs: 1) Bar graphs showing the change in relative transcript levels (“Log2FC”, log_2_ of the ratio of transcript levels) of genes representative for specific GO terms or other summaries of biological function; and 2) MA plots illustrating the relationship between Log2FC and mean transcript level for a population of related genes. In all cases, the transcriptome data for a single gene is based upon all overlapping transcripts from a common promoter.

#### I. Transcription

The structure and function of eukaryotic RNA polymerases has been recently reviewed [20]. The three RNA polymerases (i.e., Pol I, Pol II, and Pol III, synthesizing primarily ribosomal RNA, protein coding mRNA, and tRNA, respectively) possess both specific and common protein subunits. Our transcriptome data indicates that multiple subunits for each polymerase, including all of the common subunit genes, increase relative transcript levels during acute sucrose stress (Fig 3 and S5 Table). Combined with Fig 2, which establishes the persistence of activated RNA Pol II in osmotically stressed cells and Table 2, which demonstrates that relative transcript levels of many genes increase between 30 and 60 minutes, our data supports the hypothesis that RNA polymerase activity continues (perhaps more slowly) during acute hyperosmotic conditions.

**Fig 3.**
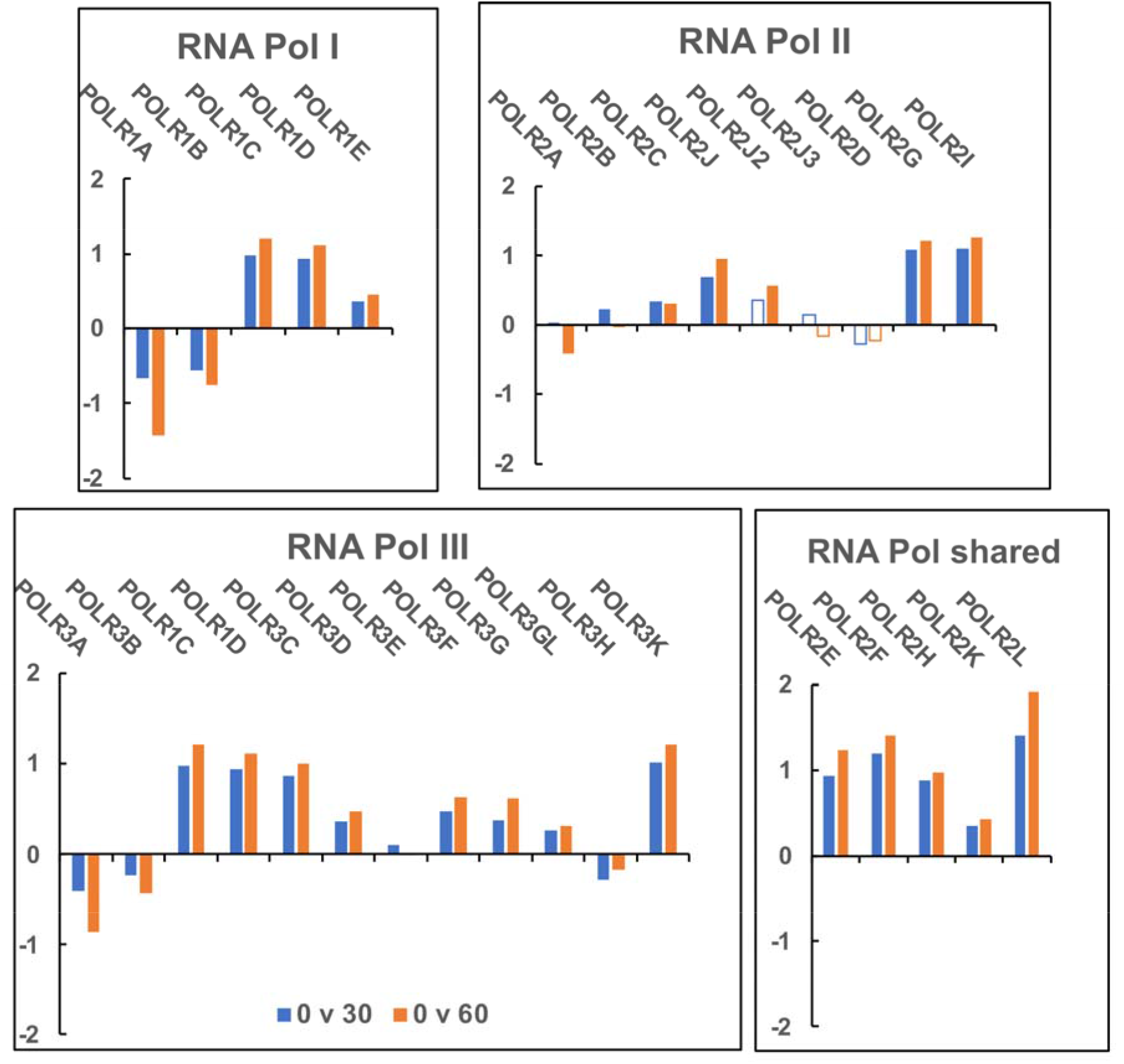
Relative transcript levels of RNA Pol I, II and III protein subunit genes in HL-60/S4 cells exposed to tissue culture medium (+300 mM sucrose) for 30 and 60 minutes. Polymerase-specific subunit transcripts are displayed in individual panels, with shared subunit transcripts in a separate panel. See S5 Table for complete data. Y-axis: Log2FC, log_2_ of the ratio of transcript levels between the hyperosmotic stress and control conditions. Solid bars signify that the change in transcript level is significant (PPDE>0.95). Open bars signify that the change in transcript level is not significant (PPDE<0.95). Sucrose exposure times: 30 min (blue); 60 min (orange). Gene codes are displayed above the relative transcript level bars.

#### II. Translation

The GO term “ribosome” (GO:0005840) includes proteins involved in ribosome assembly/disassembly, localization, binding, as well as cytosolic and mitochondrial ribosome structure. Of the 205 mapped genes in this term, 86% have significantly increased transcript levels after 30 minutes of exposure to sucrose (Fig 4a). Among the 10 genes with significantly decreased transcript levels are EIF2AK2, EIF2AK3, and EIF2AK4, which are kinases that phosphorylate translation initiation factor EIF2A to <u>inhibit</u> translation. Downregulation of these genes is consistent with increased translation. Transcript levels of all of the genes encoding the structural proteins of the cytosolic ribosome [21] increase by a factor of about 2.5x and 3.3x after 30 and 60 minutes of exposure, respectively (Fig 4b). In addition, of the 271 mapped genes in the GO term “ribosome biogenesis” (GO:0042254), 62% have significantly increased transcript levels after 30 minutes of exposure to sucrose (Fig 4c). Given that it takes only a few minutes to translate the average-size protein [22, 23], we would expect that the cellular physiological state is changing continuously during the hyperosmotic stress.

**Fig 4.**
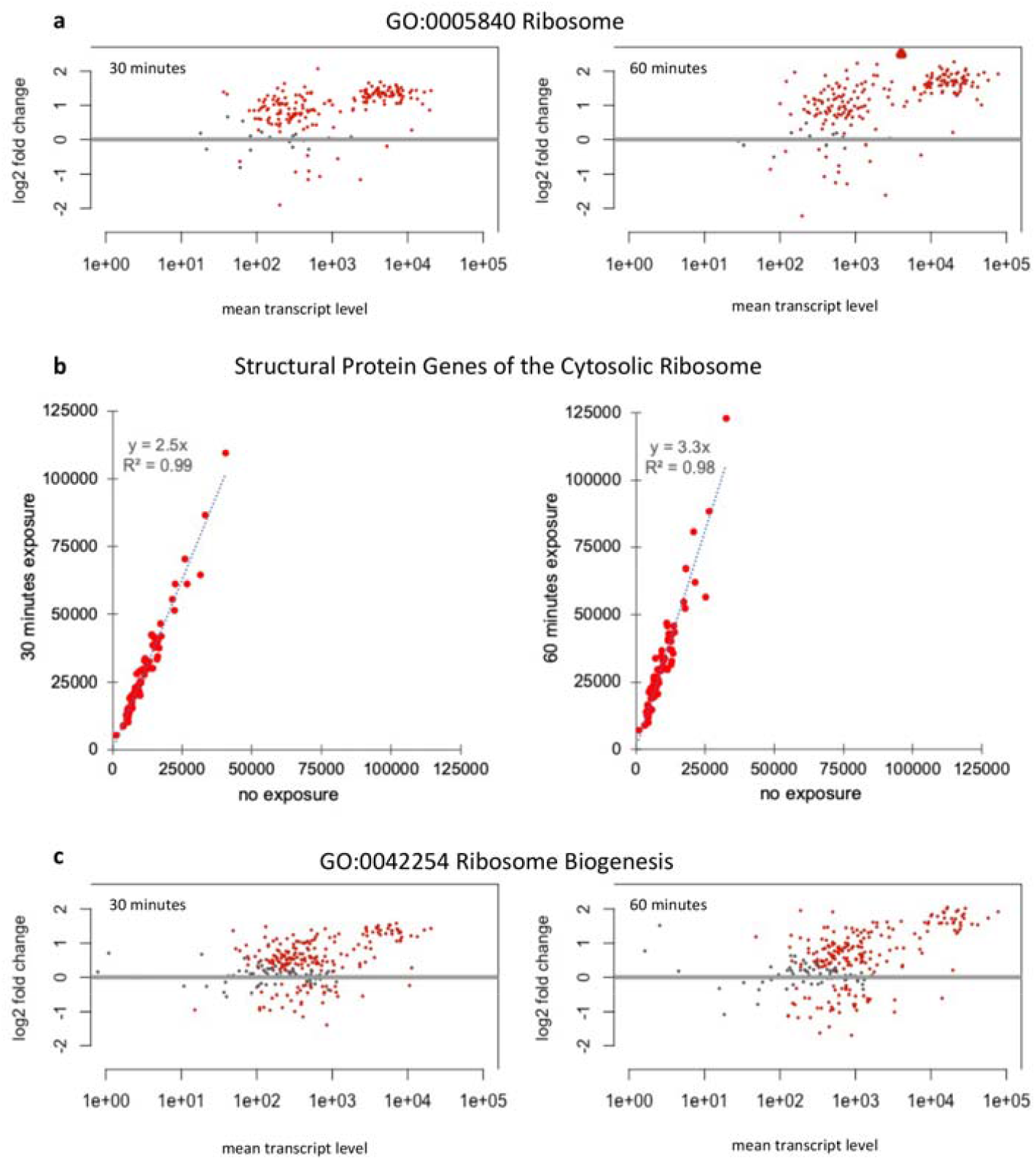
Relative transcript levels of ribosomal protein genes exposed to hyperosmotic stress. For all panels, a dot represents one gene: a red dot indicates a significant change in transcript level (PPDE>0.95); a grey dot indicates a non-significant change (PPDE<0.95). Genes with log2FC values outside the range of the Y-axis are indicated by a red arrowhead. **a)** MA plots of all mapped genes associated with GO term “Ribosome” (GO:0005840) after 30 and 60 minutes of exposure to sucrose. The X-axis is the mean transcript level of exposed and unexposed (control) conditions. The Y-axis is log2FC of the ratio of transcript levels from exposed and unexposed (control) conditions. **b)** Scatterplots of transcript levels of structural protein genes of the cytosolic ribosome with linear regression lines demonstrating that despite relative differences in transcript levels across genes, all genes increase transcript levels by about 2.5x after 30 minutes and 3.3x after 60 minutes. **c)** MA plots of all mapped genes associated with GO term “Ribosome Biogenesis” (GO:0042254) after 30 and 60 minutes of exposure to sucrose.

#### III. Mitochondria and Oxidative Phosphorylation

High concentrations of NaCl are known to cause depolarization of the mitochondrial membrane in a variety of cell types [2]. In our earlier study on the effects of acute sucrose stress upon undifferentiated HL-60/S4 cells [7], we determined that the mitochondrial membrane polarization is essentially normal for up to 1 hour of stress. Table 2 and S2 Table Biol., S3 Table Cell. and S4 Table Mol. indicate that many GO terms associated with mitochondrial structure and function are enriched in genes with increased transcript levels in cells exposed to sucrose (Fig 5). These results suggest that acute dehydration-stressed cells are attempting to maintain ATP synthesis. It is of interest to note that desiccation tolerant plants and algae are reported to increase nuclear transcription of ribosomal protein genes [24-29] and mitochondrial protein genes [24, 25], supporting a generalization of these responses in plants and animals.

**Fig 5.**
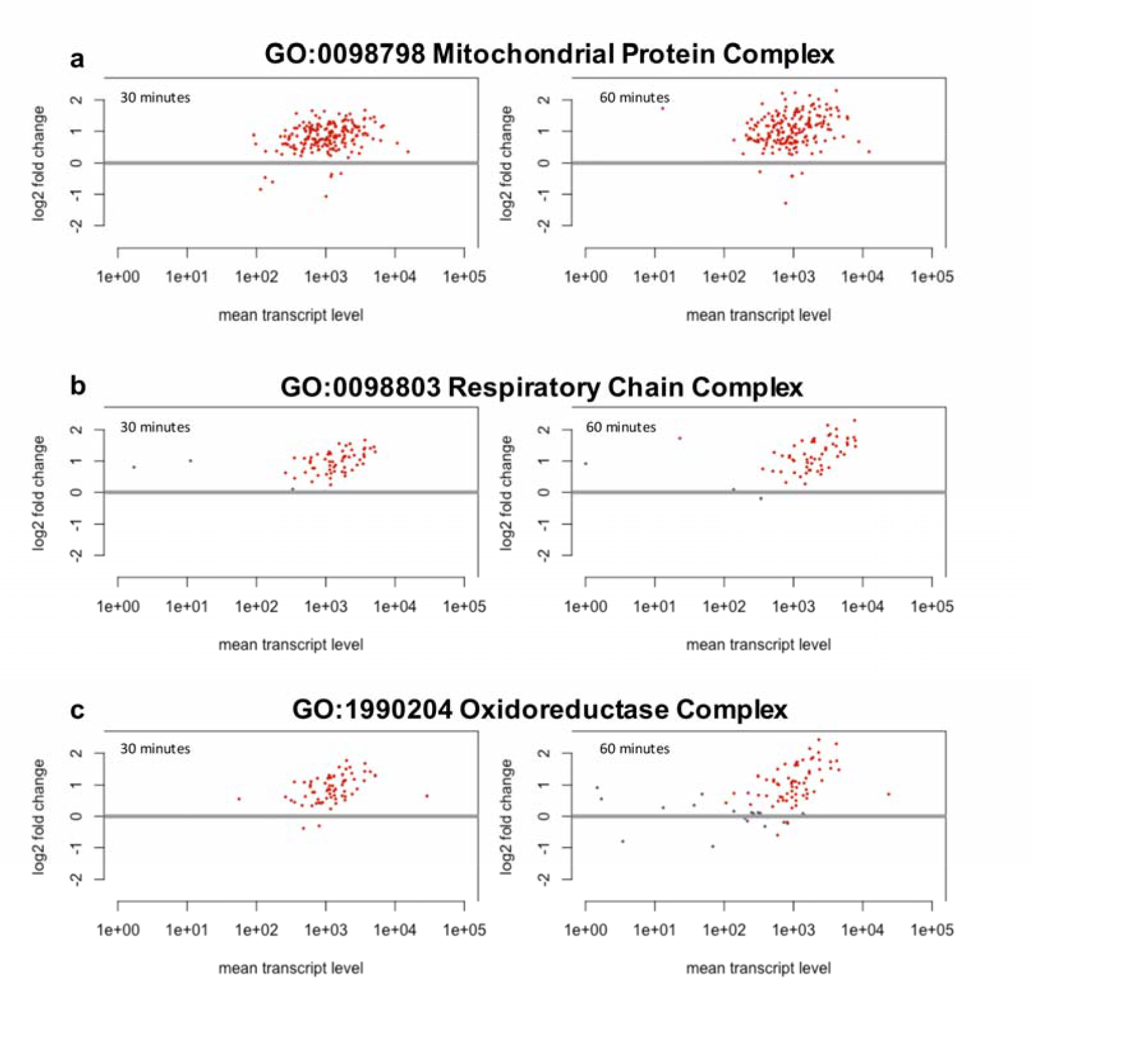
Relative transcript levels of mitochondrial protein genes exposed to hyperosmotic stress. All panels show MA plots where the X-axis is the mean transcript level of exposed and unexposed control conditions, and the Y-axis is log2FC of the ratio of transcript levels from exposed and unexposed control conditions. Panels: Left, 30 minutes of exposure; Right, 60 minutes. Each dot represents one gene: red indicates a significant change in transcript level (PPDE >0.95); grey indicates a non-significant change (PPDE<0.95). **a)** Of the 230 genes in the Mitochondrial Protein Complex, 80% exhibit significantly increased relative transcript levels after 30 or 60 minutes of exposure. **b)** Of the 60 mapped genes in the Respiratory Chain Complex, 95% exhibit significantly increased relative transcript levels after 30 or 60 minutes of exposure. **c)** Of the 90 mapped genes in the Oxidoreductase Complex, 70% exhibit significantly increased relative transcript levels after 30 or 60 minutes of exposure.

#### IV. Proteasome Activity

The GO term “proteasomal protein catabolic process” (GO:0010498) is enriched in both genes with increased and genes with decreased transcript levels. Genes with increased transcript levels include nearly all those encoding the proteasome. In addition, ubiquitin genes UBB, UBC, UBA52, and RSPS27A, as well as three E1 ubiquitin-activating enzymes, 13 E2 ubiquitin-conjugating enzymes, and components of the APC/C and ECS E3 ubiquitin ligase complexes have increased transcript levels. In contrast, genes encoding nine other E2 ubiquitin-conjugating enzymes and components of HCET, U-box, Cullin-Rbx, and single-RING finger E3 ubiquitin ligase complexes have decreased transcript levels. Together these results suggest an increase in proteasomal protein degradation mediated by increases and decreases in specific ubiquitination pathways (Table 2 Biological Process).

#### V. Replication-Dependent histone mRNA

After 30 minutes of acute exposure to sucrose only 2% of genes have a log_2_FC>1 increase in transcript levels; by 60 minutes of exposure this increases to 4.5%. Among those transcript levels with a log_2_FC>1 increase, 15% encode replication-dependent (RD) histone genes. In fact, all the major classes of RD histones (H1, H2A, H2B, H3 and H4) showed significantly increased transcript levels in hyperosmotically-stressed HL-60/S4 cells (Fig 6). There are multiple copies (isoforms) for each of these histone classes arranged in clusters [30], with the largest cluster, HIST1, on chromosome 6 (6p21-6p22). A minor number of isoform copies are located on chromosome 1 in three clusters (HIST2, HIST3 and HIST4); only HIST1 and HIST2 clusters are significantly expressed in our data. Fig 6 displays the significant (PPDE>0.95) increased levels of histone class transcripts, combining RNA-seq reads for all isoforms for each histone gene class from both clusters HIST1 and HIST2. A complete listing of transcript level changes for the histone gene isoforms is shown in S6 Table.

**Fig 6.**
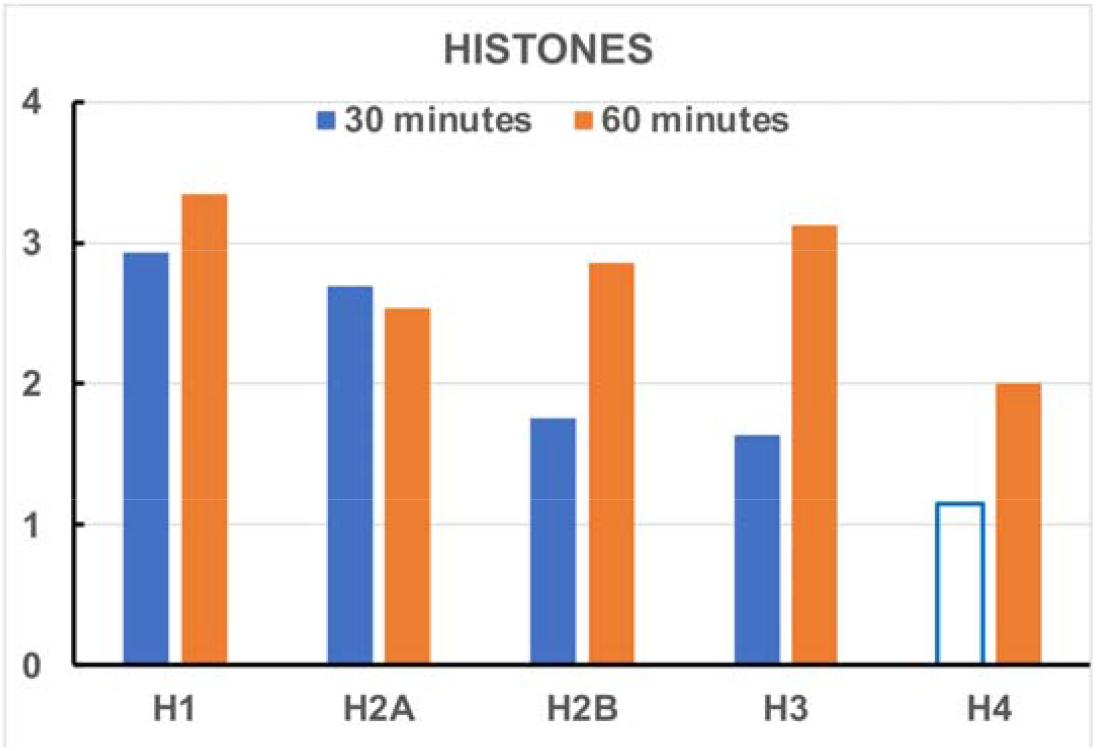
Differential expression of polyA mRNA levels of the replication-dependent (RD) histone gene classes following acute hyperosmotic stress. Only isoforms with statistically significant changes (PPDE>0.95) were included for each gene class. Y-axis: Log2FC between the control and hyperosmotic stress condition for all isoforms of each histone class after 30 (blue) and 60 minutes (orange). At 30 minutes of sucrose exposure, no histone H4 isoforms exhibited significantly increased transcript levels; at 60 minutes, one H4 isoform was significantly increased. See S6 Table for a complete list of transcript level changes for the RD histone gene isoforms.

The increase of RD histone transcript levels during hyperosmotic dehydration stress is surprising for two reasons: 1) As previously determined by cell cycle analyses [7], during this brief exposure to sucrose the % of cells in S phase was effectively unchanged (e.g., 0 min, 28.9%; 30 min, 29.9%; 60 min, 29.2%); 2) RD histone mRNAs normally have stem-loops, <u>not</u> 3’polyA tails [31, 32]. Our method of mRNA purification selects for mRNAs <u>with</u> polyA tails. Why are these stressed and apparently quiescent cells showing an <u>increase</u> in RD histone mRNA transcript levels? One plausible explanation is that the Histone Locus Body (HLB) is not functioning normally in hyperosmotically stressed cells, allowing RD histone mRNA with stem-loops to undergo a “default” conversion to 3’polyA mRNA, resulting in an apparent increase in transcript levels. The function of many of these HLB component proteins are well understood [33-35]; for example, SLBP (Stem-Loop Binding Protein) is essential for stem-loop protection at all stages of histone mRNA metabolism and NPAT (Nuclear Protein, Coactivator of Histone Transcription) is a required scaffold for HLB formation. Transcript levels of SLBP and NPAT are significantly decreased during exposure to sucrose (see S1 Table), supporting the hypothesis that the HLBs are not structurally or functionally normal. Immunostaining for NPAT (Fig 7) in cells fixed in sucrose and cells fixed in isosmotic buffer indicates that: a) HLBs are maintained during hyperosmotic stress but are more weakly stained; b) The number of HLBs/cell is lower in stressed cells. Unstressed (control) cells with functional HLBs convert only a small % of RD histone stem-loop RNA to 3’ polyA mRNA (S6 Table), see also [36], generating trace amounts of RNA-Seq reads mapping to RD histone genes, compared to the stressed cells with presumptive “disabled” HLBs. Furthermore, evidence has been published that <u>slow</u> transcription (a conceivable consequence of hyperosmotic stress) prevents formation and stabilization of the stem-loop, leading to polyadenylation [35]. We suggest that the apparent increase in RD histone transcripts during hyperosmotic sucrose stress may represent a perturbation of “normal” low-level stem-loop to polyA conversion, due to resultant changes in the composition and/or functioning of HLBs.

**Fig 7.**
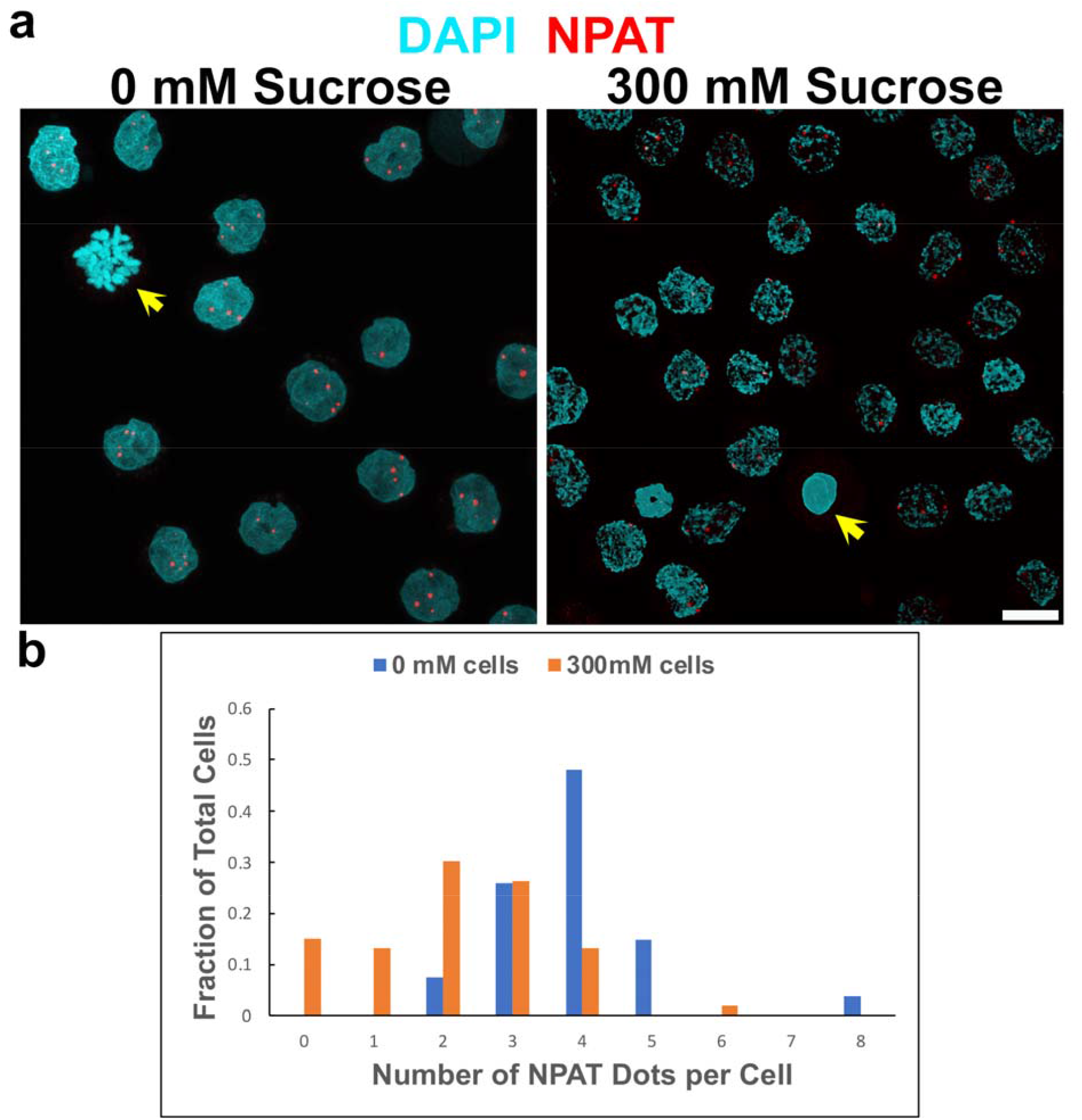
Immunostaining of NPAT, a major component of the Histone Locus Body (HLB) before and after acute hyperosmotic stress for 30 minutes. **a)** Micrographs of HL-60/S4 cells stained with anti-NPAT (red spots) within interphase nuclei stained with DAPI (cyan). Note that the mitotic chromosomes (yellow arrowheads) are not stained, and that the NPAT spots appear to be generally smaller and weaker stained in 300 mM sucrose, compared to 0 mM sucrose. **b)** Measured fraction of cells with discrete numbers of NPAT spots per cell in 0-or 300-mM sucrose. The mode values of HLBs/cell are: 0 mM sucrose, 4 HLB spots/cell (blue); 300 mM sucrose, 2 HLB spots/cell (red).

#### VI. Reversing Chromatin Repression

Acute hyperosmotic stress appears to result in relief from transcription repression, as genes with decreased transcript levels are over-represented in two GO terms involved in repression of transcription: “PcG (Polycomb Gene) Protein Complex” (GO:0031519), and “Transcription Repressor Complex” (GO:0017053). The Polycomb Gene Protein Complex includes three major groups, with complicated and overlapping interactomes [37] involved in epigenetic repression of transcription through histone modification. Key components of complexes in each group have decreased transcript levels (S7 Graph); of the 52 mapped genes associated with the GO term “Transcription Repressor Complex”, 27 experience a significant decrease in transcript levels following 30 minutes (28 genes after 60 minutes) exposure to sucrose. Collectively, these results suggest that transcription repressor effects are weakened or eliminated, consistent with the observed increase in transcript levels of many genes.

#### VII. Chromosome and chromatin structure

As documented in an earlier publication [7], protein components of the cohesin complex (e.g., RAD21 and CTCF) examined by immunostaining, appear to separate from chromatin following exposure to 300 mM sucrose. This repositioning is expected to affect the structure of TADs (“Topologically Associated Domains”, closed chromatin loops consisting of self-interacting genomic regions [38, 39], with possible effects upon gene expression. During examination of the transcriptome changes in hyperosmotic sucrose, we observed that many of the components of the cohesin ring structure exhibit significantly decreased transcript levels, relative to the transcriptome of unstressed cells (Fig 8a). This observation argues that regulation of gene expression becomes increasingly aberrant during cell dehydration.

**Fig 8.**
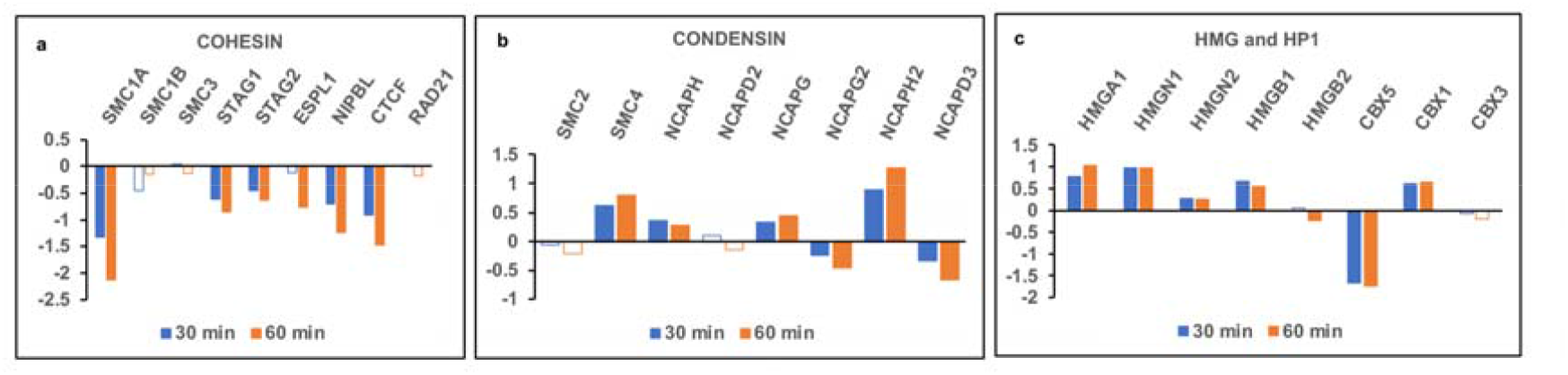
Differential polyA mRNA levels of transcripts of chromatin structural protein genes following acute hyperosmotic stress. Y-axis: Log2FC between the control and hyperosmotic stress condition at 30 (blue) and 60 minutes (red). Gene codes are displayed above the relative transcript level bars. **a)** Cohesin protein genes. HGNC Name (Common Name): ESPL1 (Separase); NIPBL (Cohesin Loading Factor). **b)** Condensin protein genes. SMC2 and SMC4 are common subunits for both Condensin I and II. **c)** HMG and HP1 protein genes. HGNC Name (Common Name): HMGN1 (HMG14); HMGN2 (HMG17); HMGB1 (HMG1); HMGB2 (HMG2); CBX5 (HP1alpha); CBX1 (HP1beta); CBX3 (HP1gamma).

Condensin I and II also build ring structures around chromatin fibers [40, 41]. Condensin I localizes on mitotic chromosomes; condensin II appears to be more important in organizing interphase chromatin. Loss of condensin can also affect gene expression, as well as centromere architecture. The actual consequences likely depend upon the chromatin context (e.g., whether the genes are active or inactive and what other proteins are bound). The consequences of changes in transcript levels in Condensin I or II to chromatin structure, during acute hyperosmotic stress, are not clear (Fig 8b).

Interphase and mitotic chromatin congelation [7] may be related to changing levels of condensins; but there is no evidence on this issue. SMC2 is a subunit component to both Condensin I and II. Immunostaining with anti-SMC2 demonstrates that it is part of an “axis” along the midline of mitotic chromosomes in normal isosmotic medium; but appears excluded from congealed mitotic chromosomes in 300 mM sucrose (See Figs 5i and j in [7]).

Exploring relative transcript levels of various well-studied chromatin binding proteins (e.g., HMG and HP1) also indicates that their transcript levels change in response to acute hyperosmotic stress in directions consistent with decreased constraints on RNA Pol II transcription (Fig 8c). Both groups of proteins can bind to nucleosomes and chromatin. To some extent, they have opposite functional consequences: HMGN1 and HMGN2 colocalize with epigenetic marks of active chromatin and with cell-type specific enhancers [42], counteracting H1 stabilization of chromatin higher-order structure [43]. HP1 proteins interact with histone tails and promote heterochromatin formation by phase separation [44, 45]. The simultaneous increase of HMGN protein transcript levels with a decrease in HP1 protein transcript levels argues for a relaxation of repression of gene expression during acute hyperosmotic stress.

## Discussion

Hyperosmotic stress in the human body is associated with inflammation and disease [1]. In addition, many healthy tissues (e.g., kidneys, liver, lymphoid tissues, the cornea, gastrointestinal tract, intervertebral discs and cartilage in joints) experience transient hyperosmotic stress during normal functioning [1]. Blood cells circulating around and through these tissues should also experience acute osmotic changes. The renal (kidney) environment experiences very significant osmotic stress. For example, during the course of a normal night’s sleep (without water intake) urine can concentrate up to ∼4x iso-osmotic “plasma” conditions [46]. Tissue-specific resilient responses allow cells to survive these osmotic stresses, but there have been few studies of genome-wide changes in gene transcription and, as far as we are aware, none involving sucrose and leukocytes.

The purpose of the present study was to gain some understanding of how the genome of an *in vivo* human cell responds to acute (i.e., 30 and 60 minutes at ∼600 milli-osmolar) stress. In our model cellular system, we exposed rapidly growing undifferentiated human leukemic promyelocytes (HL-60/S4) to acute hyperosmotic stress (medium+300 mM sucrose, ∼600 mOsM). These dehydrating conditions lead to rapid cell shrinkage, and interphase and mitotic chromatin congelation with apparent phase separation, including loss of colocalization (i.e., presumptive “demixing”) of various chromatin-associated proteins [7]. The cells remain alive and apparently healthy for at least one hour, although longer exposure results in cell death within about 24 hours. Clearly complex biophysical changes are occurring within the shrinking cells, crowding nuclei, and in the chemical environment surrounding nuclear chromatin [47]. We employed mRNA transcriptome analysis as an indicator of consequential genomic functional responses. Transcript levels at any time during the hyperosmotic stress represent the dynamic balance of mRNA transcription and degradation. It is of interest that the transcript levels of major mRNA degradation factors and enzymes are significantly reduced (e.g., log2FC approximately -1 or lower) during the hyperosmotic stress: XRN1, the main mRNA 5’-3’ exonuclease; CNOT1, deadenylation; DCP2, decapping. See S1 Table for the log2FC values after 30 and 60 minutes of dehydration. The <u>specific</u> changes in mRNA transcript levels that we observe are difficult to explain in the face of an apparent <u>general</u> decrease in mRNA degradation factors. Recent publications [48, 49] address a complex feedback mechanism between mRNA degradation and transcription. Clearly, similar research approaches are needed in order to understand the dehydration effects upon transcript levels in the HL-60/S4 cell system.

Acute hyperosmotically-stressed undifferentiated HL-60/S4 cells exhibited differentially <u>increased transcript levels</u> for genes involved in transcription, translation, and mitochondrial function, as if the cells are embarking upon active growth. Increased transcript levels for proteosome functions also occurs, suggesting accelerated protein turnover. The most surprising observation was the increase in transcript levels of replication-dependent (RD) histones. This was unexpected because the RD histone transcripts normally possess 3’stem-loops, rather than polyA tracks (which was the basis for mRNA purification). Our speculation is that the histone locus body (HLB), which protects the stem-loops during S-Phase, is not functioning properly in the hyperosmotically stressed cells. These presumptive malfunctioning HLBs may permit increased conversion of the stem-loops to 3’ polyA mRNA, producing an apparent increase in RD histone transcripts.

Genes with decreased transcript levels were enriched in various GO terms involved in repression of transcription: e.g. (GO:0031519) “PcG (Polycomb Gene) Protein Complex” and (GO:0017053) “Transcription Repressor Complex”), which might indicate more permissive conditions for mRNA synthesis (See S2 Table Biol., S3 Table Cell. and S4 Table Mol).

Other acute transcript level changes were seen in genes involved with 1) controlling chromatin domain structure (e.g., cohesin and condensin); 2) relaxing transcription repression and destabilizing heterochromatin (i.e., HMG and HP1 proteins).

Beyond the broad GO interpretations described above, the stressed cell physiology <u>cannot</u> be described as a stable state, especially in comparison to our previous analysis of chemically induced HL-60/S4 differentiation (for 4 days) into granulocytes and macrophage [12]. In that situation, the more stable phenotypic properties of the induced granulocyte and macrophage cell states were readily observed (microscopically) and in reasonable agreement with the corresponding transcriptome data. In contrast, the dehydrated HL-60/S4 transcriptome changes suggest (to us) a cellular attempt to build and grow, in the face of inevitable death. The future mode of cell death is not readily apparent in the transcriptomes. Examination for evidence of oncoming apoptosis is conflicting (e.g., transcript levels were significantly <u>decreased</u> for both “pro-apoptotic” Initiator and Effector caspases [CASP2 and CASP3] <u>and</u> for “anti-apoptotic” BCL2). Autophagy and necrosis also did not provide incriminating transcript changes. Nor is it evident that an “early osmotic stress response” [50] is occurring following the acute hyperosmotic stress. The relative transcript levels for early response genes NFAT5 (TonEBP) and SLC6A6 (TauT) both <u>decrease</u> sharply at 30 and 60 min. One clue suggests that oxidative stress may play a role in the eventual cell death. Oxidative stress can occur when mitochondria produce more reactive oxygen species (ROS) than available antioxidant defenses can mitigate, resulting in critical cell damage. The transcript levels of many genes involved in glutathione-and thioredoxin-based antioxidant defense are elevated at 30 and 60 minutes of hyperosmotic stress, suggesting that ROS are increasing. In particular, TXN2 (thioredoxin 2) protects against oxidative stress in mitochondria, inducing cells to become insensitive to ROS-induced apoptosis [51]. If TXN2 transcript levels should decrease after 60 minutes exposure, or if the total amount of thioredoxin 2 (and other antioxidant proteins) are unable to compensate for the levels of ROS produced by hyperosmotic stress, ROS-induced apoptosis could be fatal [52].

Despite this incomplete understanding of the physiological state of the hyperosmotically-stressed cells, there is compelling evidence that the structure of chromatin is profoundly altered. This is readily apparent in the earlier immunostaining study [7], which indicated an apparent phase separation and demixing of cohesin (RAD21) and condensin (SMC2) components, CTCF, histones H1.2 and H1.5, and non-histones HMGN2 and HMGB2. A previous study [53] employing hypertonic NaCl (which is also hyperosmotic) on breast cancer T47D cells analyzed chromatin changes using Hi-C and ChIP-seq (i.e., the DNA sites of bound RNA Pol II, CTCF and RAD21). Hypertonic stress resulted in a decrease in the number of TADs, a weakening of TAD boundaries and major perturbations of Compartments A and B (i.e., euchromatin and heterochromatin regions). RNA Pol II became dislocated from many transcription start sites (TSS) and appeared to run-off transcription end sites (TES). In addition, CTCF and RAD21 were displaced from their normal binding sites, consistent with our immunostained images [7]. Recent studies underscore the importance of normal cohesin function to higher-order interphase nuclear architecture [54].

The concept of Liquid-Liquid Phase Separation (LLPS) or “condensates”, where cellular macromolecules can be either concentrated into microscopic phases or dissolved in a cellular solution, depending upon their specific “critical concentration for phase separation”, has become a major biophysical perspective for interpreting non-membranous cellular particles, including chromatin [55-60]. Accepting that acute hyperosmotic stress with either NaCl [53] or sucrose [7, 61] produces profound structural changes of *in vivo* chromatin (described as “altered” phase separation), we argue that this perturbed chromatin may yield somewhat disorganized transcriptomes and aberrant cell physiological states. We suggest that, during acute dehydration, condensates may concentrate or disperse molecules in disordered or nonfunctional ways, dictated by the changing biophysical environment. Furthermore, we suggest that “unstressed” chromatin regions normally possess biophysical microheterogeneity of their “critical concentration for phase separation”. Most likely, this biophysical microheterogeneity is due to local differences in protein composition, post-translational modifications and physiologic state at the moments prior to the acute hyperosmotic shock.

Disorganized gene expression can have rapid effects. Transcription and translation of average size mRNA and protein (e.g., 500 amino acids) takes ∼1 minute for each process [22, 23]. The population of cellular proteins might exhibit significant changes by 30 and 60 minutes of osmotic stress. Decreases and/or displacement of chromatin structural proteins (e.g., components of cohesin and condensin) during acute osmotic shock could produce disruption of TADs [62], which may further exacerbate transcription disorder. From this perspective, we should not be surprised that cell differentation, which follows evolutionary determined transcription programs in a methodical manner, yields coordinated transcriptomal responses; whereas, acute hyperosmotic stress and cell dehydration results in transcriptomes that are less focused on specific phenotypes. In the present study, the broad GO functions exhibiting relatively <u>increased</u> transcript levels (e.g., transcription, translation and mitochondrial oxidative phosphorylation) suggest that stressed chromatin can still yield transcriptomes with some functional coordination. We suggest that this hyperosmotic stress response provides a useful *in vivo* model for examining rapid nuclear and chromatin biophysical changes (e.g., altered phase separation) that can influence chromatin higher-order structure and the regulation of transcription.

## Conclusion

The induced physiological state of acute hyperosmotically stressed undifferentiated HL-60/S4 cells resembles a rapid attempt to rebuild the damaged cells by increasing transcript levels of genes involved in transcription, translation, and energy production. This stressed cell physiologic state is not as well defined as the chemically-induced differentiated cell states of HL-60/S4, designated “granulocytes” and “macrophage” [12, 13]. The suddenness of hyperosmotic stress with resultant changes in solute concentrations and macromolecular crowding, very likely yields altered phase separation with somewhat uncoordinated gene expression, ultimately unable to prevent cell death. Ironically, it might be the “Raging” of the ROS that results in the final “dying of the light.”

## Supporting information

S2_Table Biol

S3_Table Cell

S4_Table Mol

S7_Graph

S8_Text run_rsem_script

## Acknowledgments

The authors express their gratitude to the Marine Biology Laboratory, to Bates College and to the School of Pharmacy, University of New England for space and support, and to Logan Boyd for assistance with STED imaging.

## Supporting Information

**S1 Table. RSEM-EBSeq results**. Log2(PostFC): log base 2 of the posterior calculation of fold change between conditions; PPDE: Posterior probability of differential expression. C1Mean: normalized mean count of reads mapped in condition 1 (control or 30 minutes exposure); C2Mean: normalized mean count of reads mapped in condition 2 (30 or 60 minutes exposure). See EBSeq manual for details. Genes with empty cells for PPDE, counts, etc. indicate that no reads mapped to those genes.

**S2 Table Biol. WebGestalt output from genes with significantly decreased transcript levels after 60 minutes**. 2270 genes mapped. Size: number of human genes in gene set; Expect: expected number of genes mapped to gene set; Ratio: number of genes mapped over expected; P value: single test significance; FDR: false discovery rate based on BH multiple correction; % GO: percent of genes in GO term mapped; % mapped: percent of all mapped genes in GO term.

**S3 Table Cell. WebGestalt output from genes with significantly decreased transcript levels after 60 minutes**. 1676 genes mapped. Size: number of human genes in gene set; Expect: expected number of genes mapped to gene set; Ratio: number of genes mapped over expected; P value: single test significance; FDR: false discovery rate based on BH multiple correction; % GO: percent of genes in GO term mapped; % mapped: percent of all mapped genes in GO term.

**S4 Table Mol. WebGestalt output from genes with significantly decreased transcript levels after 60 minutes**. 1938 genes mapped. Size: number of human genes in gene set; Expect: expected number of genes mapped to gene set; Ratio: number of genes mapped over expected; P value: single test significance; FDR: false discovery rate based on BH multiple correction; % GO: percent of genes in GO term mapped; % mapped: percent of all mapped genes in GO term.

**S5 Table. Transcript level changes in genes encoding peptides in the holoenzymes RNA Pol 1, RNA Pol 2 and RNA Pol 3 after 30- and 60-minutes exposure to sucrose**.

**S6 Table. RSEM-EBSeq results for genes encoding histone proteins**. PPDE: Posterior probability of differential expression; see EBSeq manual for details. Empty cells for PPDE etc. indicate no reads mapped to that gene. Columns S-AA show averages for each isoform from each cluster.

**S7 Graph. Changes in polyA mRNA transcript levels in Polycomb Gene Protein complexes**. Y-axis: Log2(FC) between transcript levels of the hyperosmotic stress condition and control. Blue, 30 minutes; Orange, 60 minutes; solid bars, PPDE > 0.95; open bars, PPDE < 0.95.

**S8 Text. BASH Script of the workflow**.

