## Supplementary material for "The transcriptome of acute dehydration in Myeloid Leukemia cells": S7_Graph

**S7 Graph. Changes in polyA mRNA transcript levels in Polycomb Gene Protein complexes.** Y-axis: Log2(FC) between transcript levels of the hyperosmotic stress condition and control. Blue, 30 minutes; Orange, 60 minutes; solid bars, PPDE > 0.95; open bars, PPDE < 0.95.

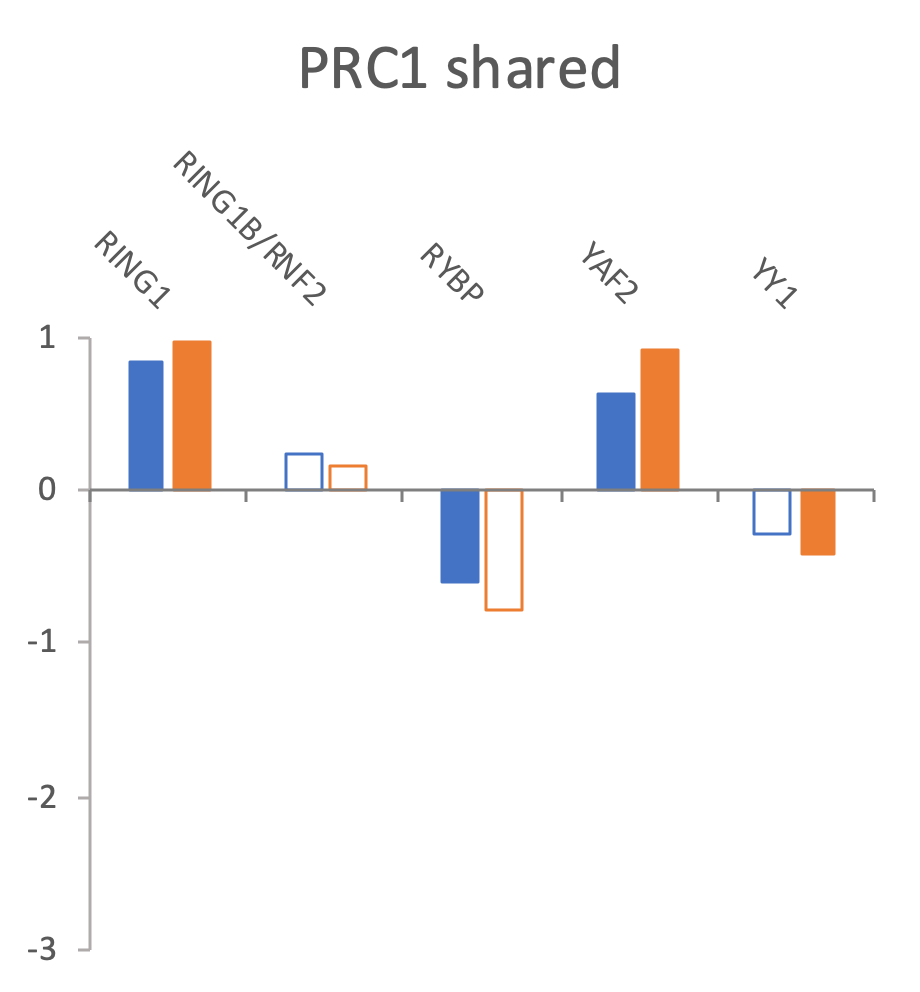

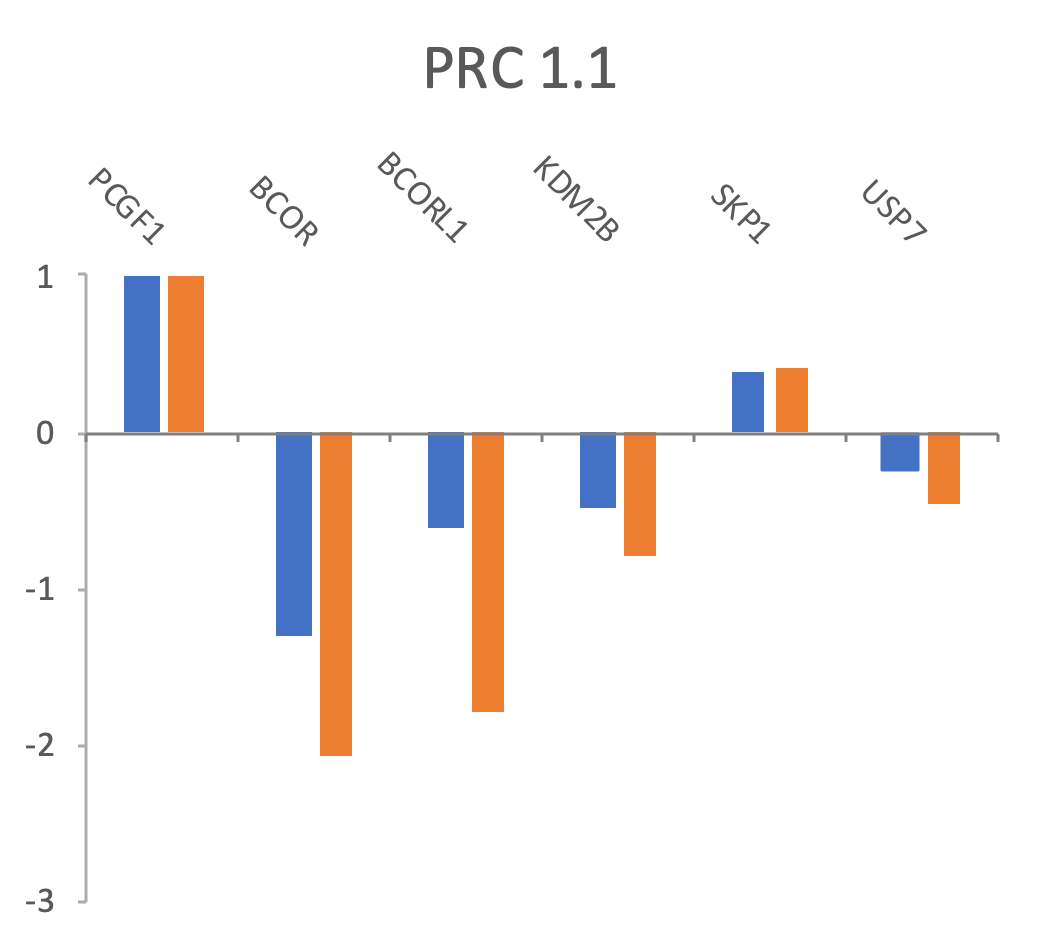

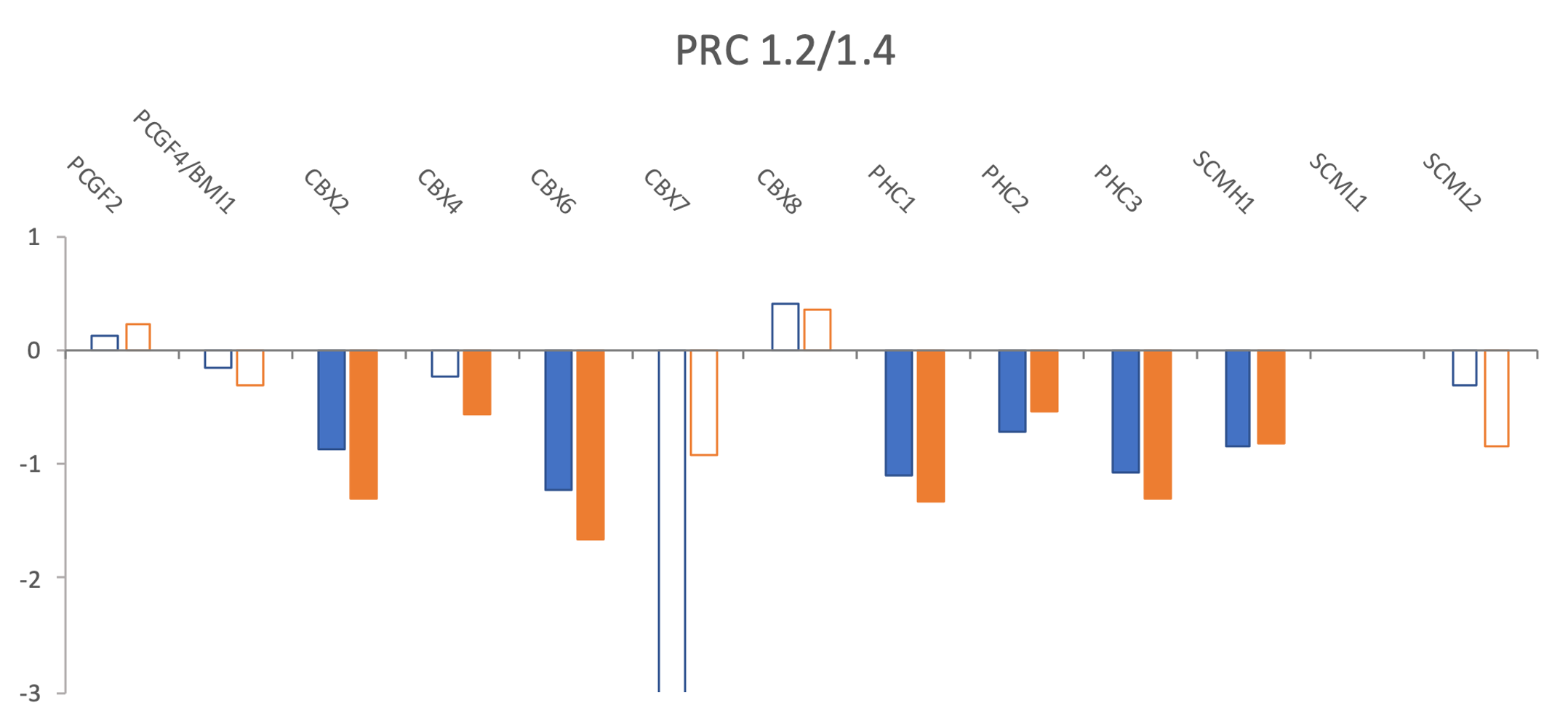

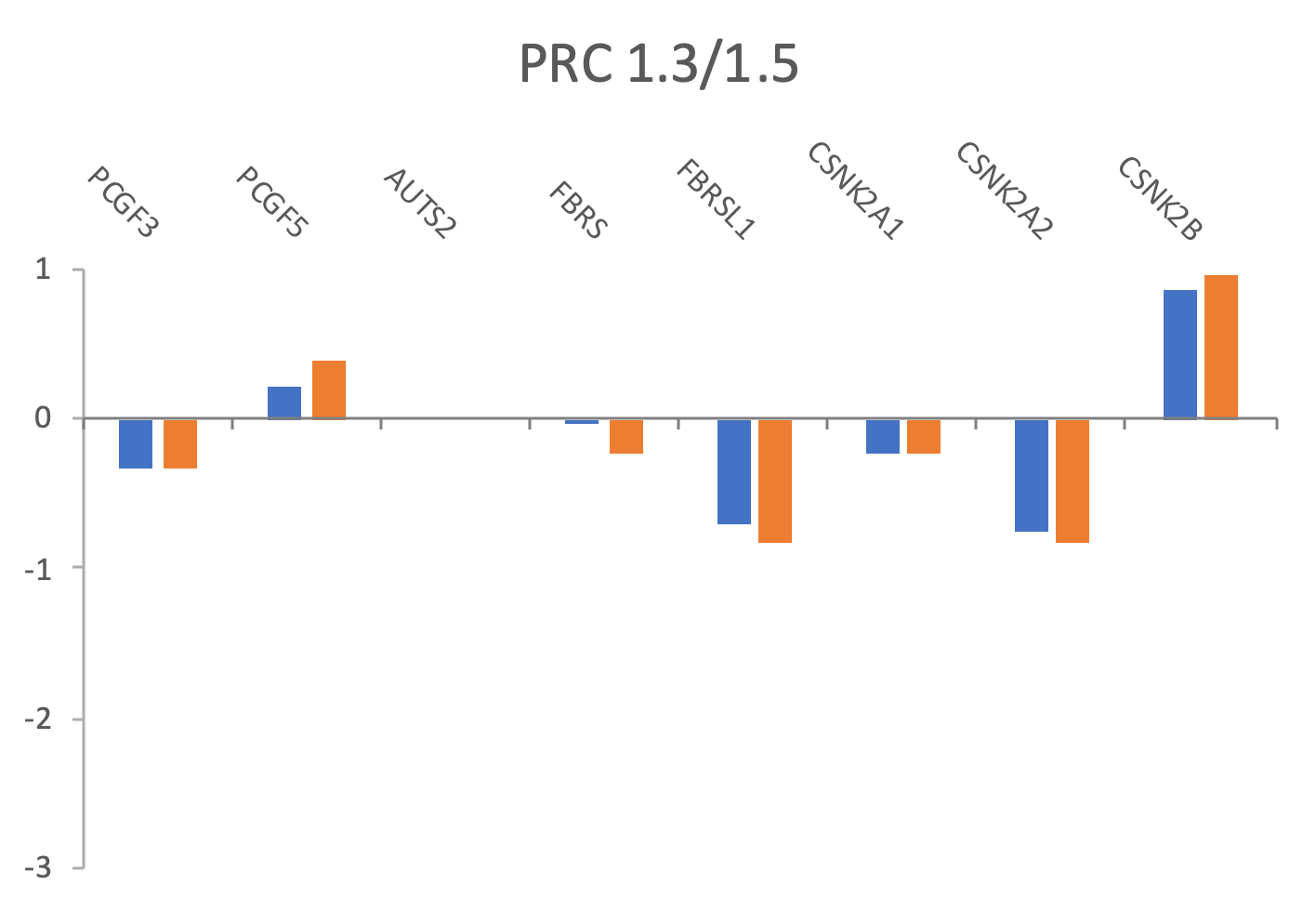

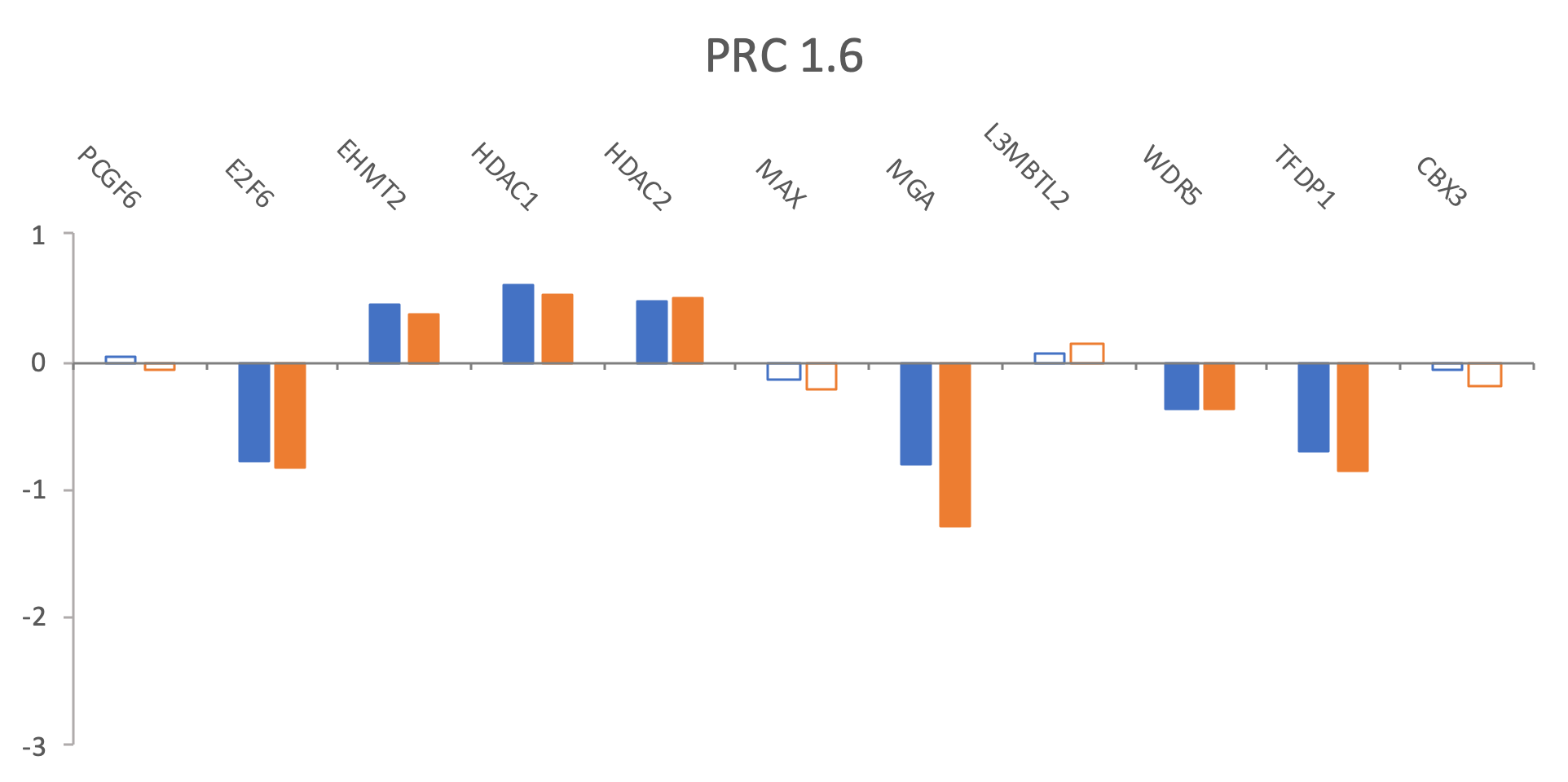

PRC1 shared

PRC1.1

PRC1.2/1.4

PRC1.3/1.5

PRC1.6

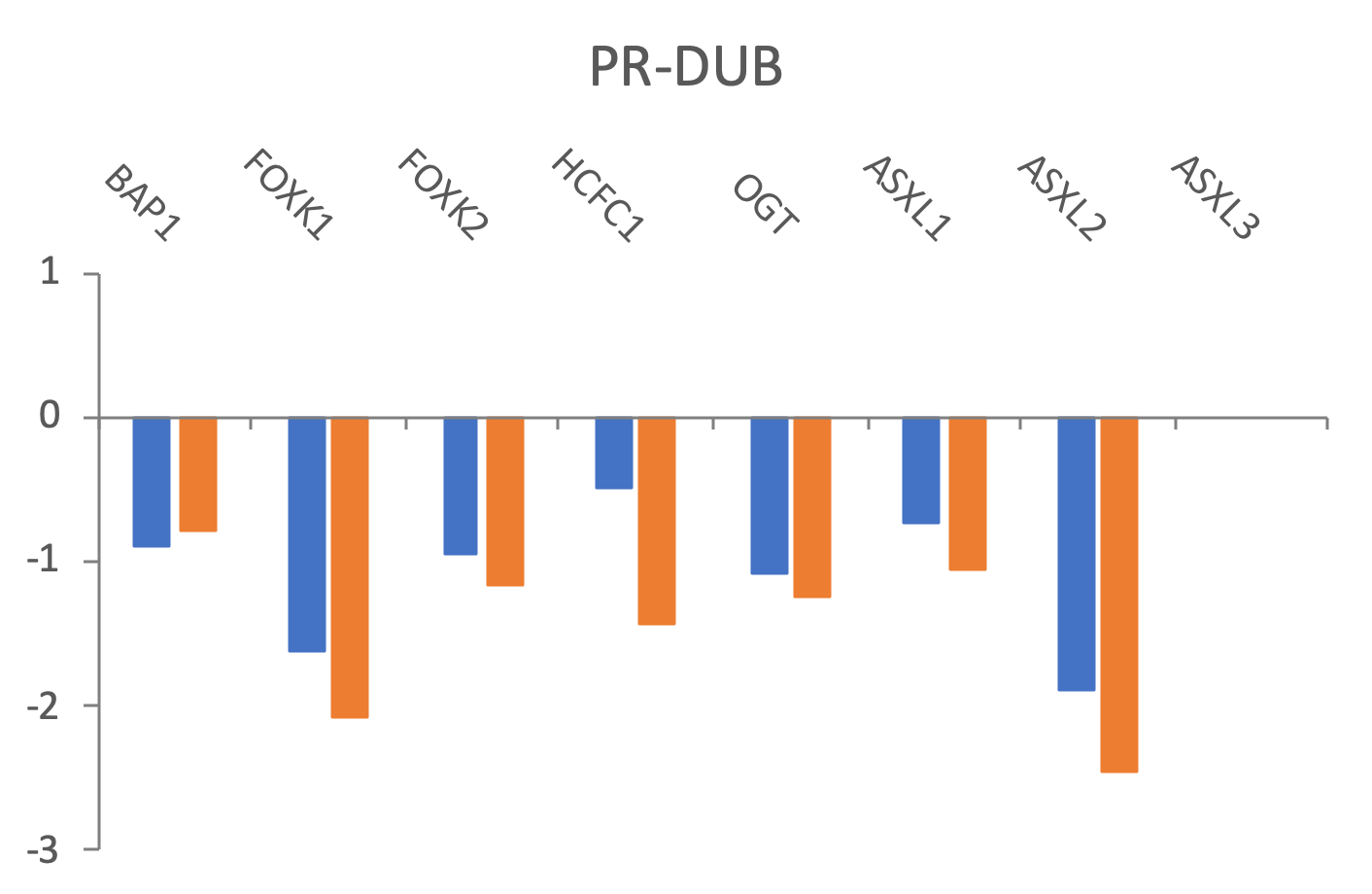

PRC2

PR-DUB

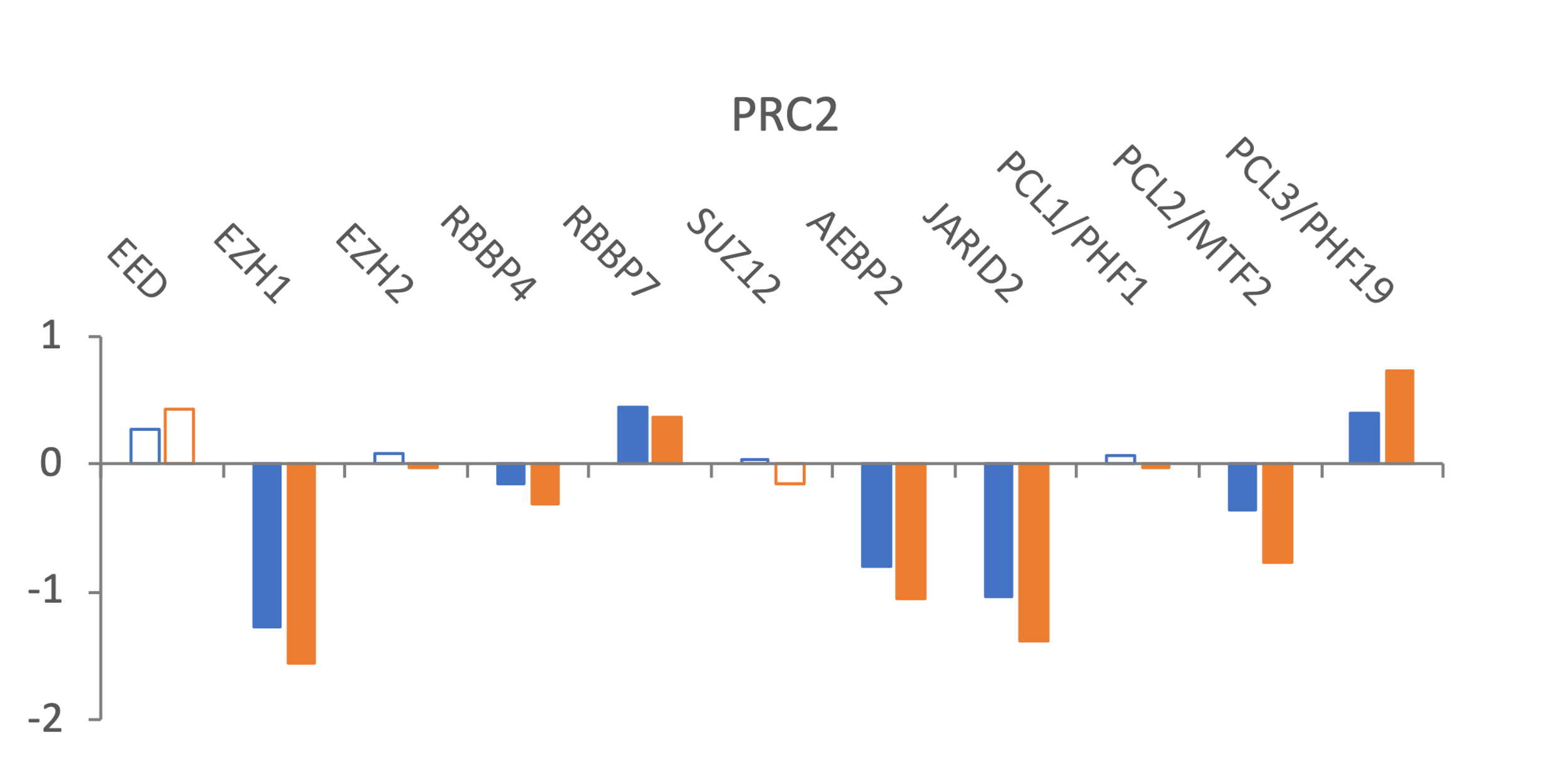

PRC2
